# Molecular Phylogeny of human adenovirus type 41 lineages

**DOI:** 10.1101/2022.05.30.493978

**Authors:** Jasper Götting, Anne K Cordes, Lars Steinbrück, Albert Heim

## Abstract

Type 41 of human adenovirus species F (HAdV-F41) is a frequent aetiology of gastroenteritis in children, and nosocomial as well as kindergarten outbreaks have been frequently described. In contrast to other HAdV types, HAdV-F41 was not associated with life-threatening disseminated disease in allogeneic haematopoietic stem cell transplant (HSCT) recipients or any severe organ infections so far. Due to the limited clinical significance, the evolution of HAdV-F41 has not been studied in detail. Recently, HAdV-F41 has been associated with severe hepatitis in young children, and interest in HAdV-F41 has skyrocketed, although the aetiology of the hepatitis has not been resolved.

Complete genomic HAdV-F41 sequences from 32 diagnostic specimens of the past 11 years (2011–2022) were generated, all originating from gastroenteritis patients. Additionally, 33 complete HAdV-F41 genomes from GenBank were added to our phylogenetic analysis.

Phylogenetic analysis of 65 genomes indicated that HAdV-F41 evolved with three lineages co-circulating. Lineage 1 included the prototype ‘Tak’ from 1973 and six isolates from 2007 to 2017 with an average nucleotide identity of 99.3 %. Lineage 2 included 53 isolates from 2000 to 2022, had an average nucleotide identity of 99.8 %, and split into two sublineages. Lineage 3, probably described for the first time in 2009, had a 45nt deletion in the long fiber gene and had evolved significantly in the short fiber and E3 region. Moreover, a recent lineage 3 isolate from 2022 had a recombinant phylogeny of the short fiber gene. Fibers interact with cellular receptors and determine cellular tropism, whereas E3 gene products interfere with the immune recognition of HAdV infected cells.

This in-depth study on the phylogeny of HAdV-F41 discovered significant evolution of recently described lineage 3 of HAdV-F41, possibly resulting in altered cellular tropism, virulence and pathophysiology.

## Introduction

Human adenoviruses (HAdV) are non-enveloped, icosahedral DNA viruses, first isolated in 1953 from human adenoidal tissue (Rowe et al., 1953; Hilleman and Werner, 1954) and belong to the *Mastadenovirus* genus. Their linear dsDNA genome is ^~^35 kb in length and encodes about 30–40 proteins (Davison et al., 2003). HAdVs are further separated into seven species (A–G) by phylogenetic criteria and subdivided into 112 types with type numbering merely related to their date of first isolation. HAdV types were initially defined by cross-neutralization (HAdV serotypes 1–51) and later by sequencing of all three major capsid proteins penton, hexon, and fiber (genotypes 52–112) (Aoki et al., 2011; Seto et al., 2011).

Only two types (40 and 41) are members of species HAdV-F. HAdV-F41, prototype strain ‘Tak’, was isolated in 1973 from the stool of a child suffering from gastroenteritis in the Netherlands (de Jong et al., 1983). HAdV-F41 is associated almost exclusively with gastroenteritis, most frequently in the toddler age and is the second most frequent cause of diarrhoea in this age group, only second to Rotavirus (Lee et al., 2020). HAdV-F41 frequently causes nosocomial outbreaks in paediatric wards and community facilities such as kindergartens, but the disease remains limited to the gut even in severely immunosuppressed children (Mattner et al., 2008; Gonçalves et al., 2011; Lefeuvre et al., 2021). Only a single case of severe HAdV-F41 dissemination has been reported in a stem cell transplant recipient (Slatter et al., 2005). This uniquely restricted organ tropism of HAdV-F41 can be attributed to the absence of an RGD motif in its penton base, which binds types of all other HAdV species to its secondary cellular receptors, α_v_β_3_ and α_v_β_5_ integrins (Wickham et al., 1993; Albinsson and Kidd, 1999). Recently, it was reported that the short fiber of HAdV-F41 binds to cells via heparan sulfate (Rajan et al., 2021), which may restrict its cellular tropism. In contrast, the long fiber of HAdV-F41 binds to the cellular CAR receptor as many other types of species HAdV-A, -C, -D, and -E do (Roelvink et al., 1998).

New types may evolve via diversifying selection (immune escape) of the neutralisation determinant (Robinson et al., 2013). Recombination between types of the same HAdV species is also an essential mechanism for the evolution of species HAdV-B, -C and -D (Robinson et al., 2013). However, with its two types, HAdV-F hardly offers many options for intertypic recombination. Nevertheless, multiple distinct strains of HAdV-F41 have been distinguished by different growth characteristics, multiple restriction enzyme polymorphisms, and reactivity with different neutralising monoclonal antibodies (de Jong et al., 1983; van der Avoort et al., 1989). Despite these early studies, the molecular phylogeny of HAdV-F41 has not yet been studied in detail because of its limited clinical significance as a mere gastroenteritis virus. Recently, HAdV-F41 DNAemia was associated with severe hepatitis in young children (Baker et al., 2022; Marsh et al., 2022).

Therefore, we sequenced the genomes and analysed the molecular phylogeny of 32 HAdV-F41 clinical isolates from 2011 to 2022, all originating from gastroenteritis cases. Moreover, 33 recently published complete genomic HAdV-F41 sequences from GenBank were included in the phylogenetic analysis.

## Materials and methods

### HAdV-F41 isolates and complete genomic sequences

HAdV-F41-positive samples (stool or cell culture supernatant from A549 cultures used for virus isolation) originating from the collection of the German national adenovirus reference laboratory were sequenced as described below. Furthermore, all 33 available complete genomic HAdV-F41 sequences from GenBank were included in the phylogenetic analysis. Sequences from (Tahmasebi et al., 2020) were excluded due to incompleteness and unusual number of SNPs.

### Ethical statement

The study only analysed viral data without patient material and thus did not require approval from the ethics committee.

### High-throughput sequencing and *de novo* assembly

DNA was extracted from 400 μl HAdV-F41 positive stool or cell culture supernatant (depending on the availability and virus load) using a Qiagen Blood Kit on a QIAcube. Library preparation was performed using the NEBNext Ultra II FS DNA Library Prep Kit for Illumina according to the manufacturer’s protocol. Final libraries were inspected on an Agilent Bioanalyzer, normalised, multiplexed, and sequenced on an Illumina MiSeq using a 600v3 Reagent Kit to generate 2×300 bp paired-end reads with an average of 1.25 million reads per sample.

*De novo* assembly was performed as previously described (Dhingra et al., 2019). Briefly, human reads were removed, viral reads were trimmed with fastp and assembled with SPAdes, which usually resulted in a single high-coverage contig constituting the entire HAdV-F41 genome. Finally, genomes were polished using Pilon (Walker et al., 2014) and genome termini were manually examined and corrected using a mapping of the reads against the HAdV-F41 reference GenBank sequence (DQ315364). The resulting HAdV-F41 genomes were annotated from the HAdV-F41 reference sequence using Geneious Prime 2020.1.2. Finally, genomes were deposited in GenBank (Accession numbers ON442312–ON442330, ON532817–ON532827).

### Phylogenetic analysis

Multiple sequence alignment of the complete sequences was carried out using MAFFT v7.450 (Katoh and Standley, 2013). Phylogenetic trees were constructed with the HAdV-A61 reference sequence as the outgroup using RAxML v8 under the GTR GAMMA model with 500 rapid bootstrapping replicates and search for the best-scoring ML tree (Stamatakis, 2014). For comparison, the complete genomic sequence phylogeny was additionally inferred using MrBayes 3.2.6 (GTR & invgamma model with 500,000 MCMC steps and 50,000 burn-in steps) and Geneious Tree Builder (Neighbour-Joining with default parameters and 1000 bootstrap replicates) (Ronquist et al., 2012). SimPlot 3.5.1 was used to generate similarity plots (SimPlots) and to perform BootScan recombination analyses using default parameters, a window size of 1000 bp (BootScan) or 1500 bp (SimPlot), and a step size of 200 bp (BootScan) or 300 bp (SimPlot) (Lole et al., 1999). TreeTime with default parameters was used to attempt inference of a time-calibrated maximum-likelihood phylogeny (Sagulenko et al., 2018). *In silico* RFLP analysis was performed in Geneious Prime 2020.1.2 (Biomatters) with the ten restriction enzymes used in the previous RFLP genome typing work (BamHI, BglI, BstEII, EcoRI, HindIII, KpnI, PstI, SacI, SmaI, XhoI) (van der Avoort et al., 1989). Complete genomic sequences were digested with each enzyme separately, calculated fragments were rounded to the nearest 100 bp length, and fragments shorter than 400 bp were discarded to match the data with the fragment patterns from figure 1 of Johansson et al. 1991. A 137-bit binary string representing the presence or absence of all 137 occurring fragments from all enzymes was compiled for all complete genomic sequences as well as the 24 described genome types from table 1 of van der Avoort et al. 1989. All phylogenetic trees were visualised and annotated in R using ggplot2 and ggtree (Wickham, 2016; Yu, 2020).

**Figure 1:**
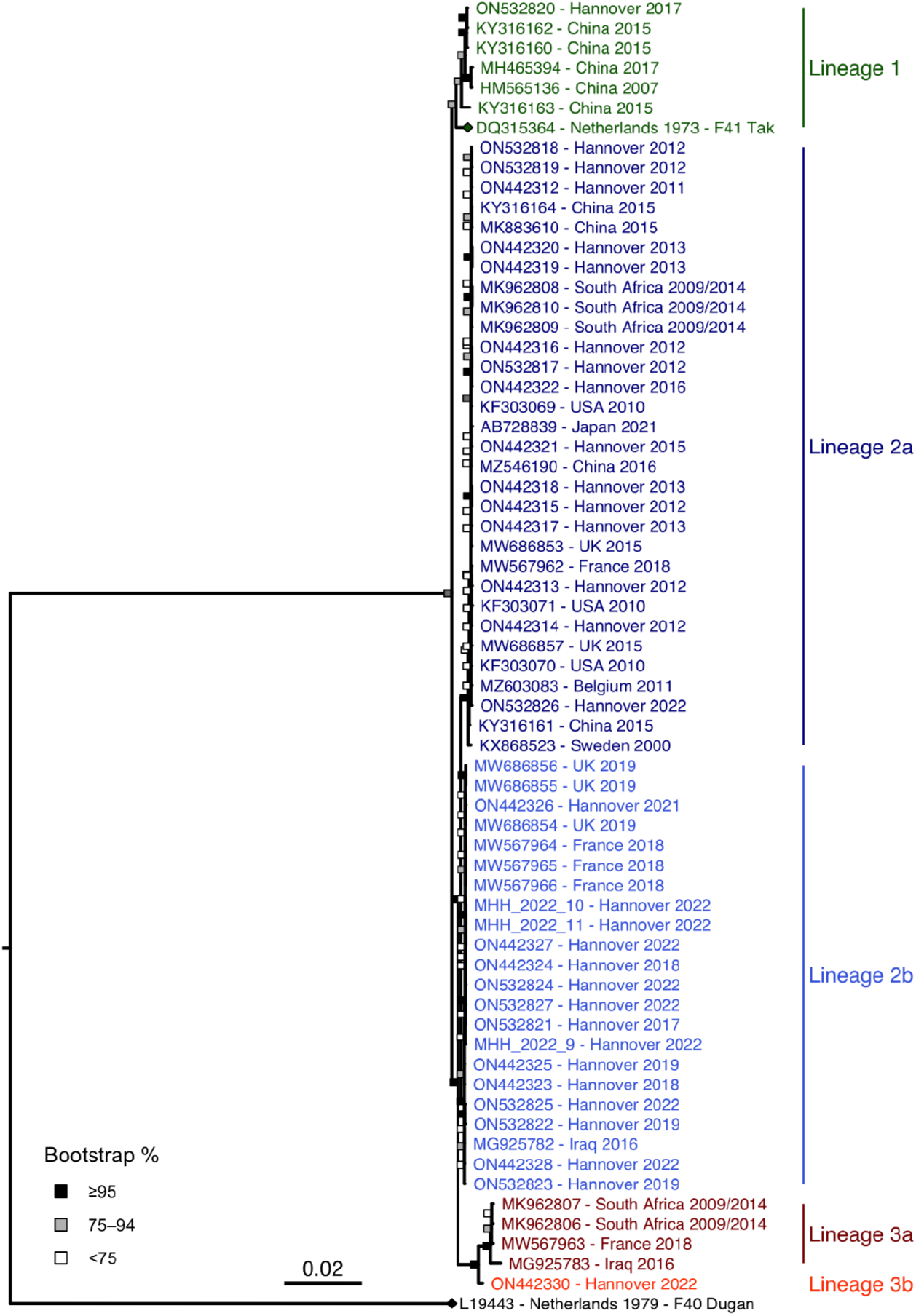
HAdV-F41 complete genomic sequence phylogeny. Maximum likelihood phylogeny of 65 HAdV-F41 genomes with the HAdV-F40 prototype as the outgroup. The two prototype sequences (HAdV-F41 DQ315364, HAdV-F40 L19443) are marked with a rhombus at the tip point. Bootstrap support values were binned into three categories (≥95 %, 75–94 %, <75 %) indicated by filled boxes at the node points.

### Analysis for positive selection

BUSTED (Branch-Site Unrestricted Statistical Test for Episodic Diversification), as implemented on datamonkey.org, was utilised to identify genes under positive or diversifying selection (Weaver et al., 2018). All genes displaying high diversity in the SimPlot were analysed in BUSTED, which tests for gene-wide, non-site-specific selection (Murrell et al., 2015).

## Results

### HAdV-F41 phylogeny

Phylogenetic analysis of 65 complete genomic HAdV-F41 sequences exhibited three distinct lineages containing multiple identical or barely divergent sequences (up to 99.9 % identity within lineages) (Figure 1). Clustering of lineages was stable between different tree models (maximum-likelihood, neighbour-joining, Bayesian inference; see Supplementary Figures 1 and 2) and confirmed by bootstrapping.

Lineage 1 included the prototype ‘Tak’ from 1973 and isolates from as late as 2017. Only seven of 65 complete genomic sequences were clustered as lineage 1, but these originated from multiple geographic regions (China, Netherlands, and Germany). Lineage 1 had a 99.3 % average nucleotide identity and even 99.2 % identity between the 1973 prototype and the last available isolate from 2017. Lineage 2, which had two sub-lineages (2a and 2b), included the majority of the genomic sequences (53 of 65), which originated from 2000 to 2022 and originated from multiple regions (Germany, Belgium, Japan, USA, Sweden, China, South Africa, France, UK, and Iraq). The nucleotide identity averaged 99.8 % within lineage 2, 99.9 % within sublineage 2a and 99.9 % in sublineage 2b. The average identity between lineages 1 and 2 was high (98.9 %), whereas lineage 3 was more divergent from lineage 1 (98.2 %). Lineage 3 included two sublineages but lineage 3b was represented only by a single sequence originating from Germany, 2022. Lineage 3a included only four sequences originating from South Africa, France and Iraq, 2009 to 2018. Nucleotide identity within lineage 3 averaged 99.6 %.

Intratypic evolution was so slow that constructing time-calibrated complete genomic sequence phylogenies was unsuccessful (Supplementary Figure 3).

### Evolution of genome regions

Hotspots of evolution separating the lineages were found in the hexon, long fiber, and the short fiber genes (Figure 2), but the penton base gene was highly conserved (99.6–100 %, Supplementary Figure 3). Only lineage 2a has evolved significantly in the hexon gene with seven non-synonymous mutations in the loops of the neutralisation determinant ε. However, the BUSTED algorithm did not confirm the positive selection of potential immune escape variants (*p*=0.087). Two lineage 3 hexon sequences did not cluster with other lineage 3 sequences, but this was not supported by significant bootstrap values. Lineage 3 had a 45-nucleotide deletion in the long fiber gene resulting in a 15 amino acids shorter fiber shaft. Furthermore, the long fiber gene of lineage 3 had 21 SNPs (9 of these non-synonymous) compared to the consensus sequence. Four of nine amino acid substitutions were located in the fiber knob which binds the cellular receptor and haemagglutination inhibiting antibodies. Evidence for positive selection was found by the BUSTED (*p*=0.000) algorithm for the entire long fiber gene but not for the knob (*p*=0.326). Only lineage 3a has evolved significantly in the short fiber gene with 51 SNPs compared to the consensus sequence resulting in 20 amino acid substitutions, with four in the knob region. However, positive selection was not confirmed by the BUSTED algorithm (*p*=0.444 for the entire short fiber gene, *p*=0.5 for the short fiber knob).

**Figure 2:**
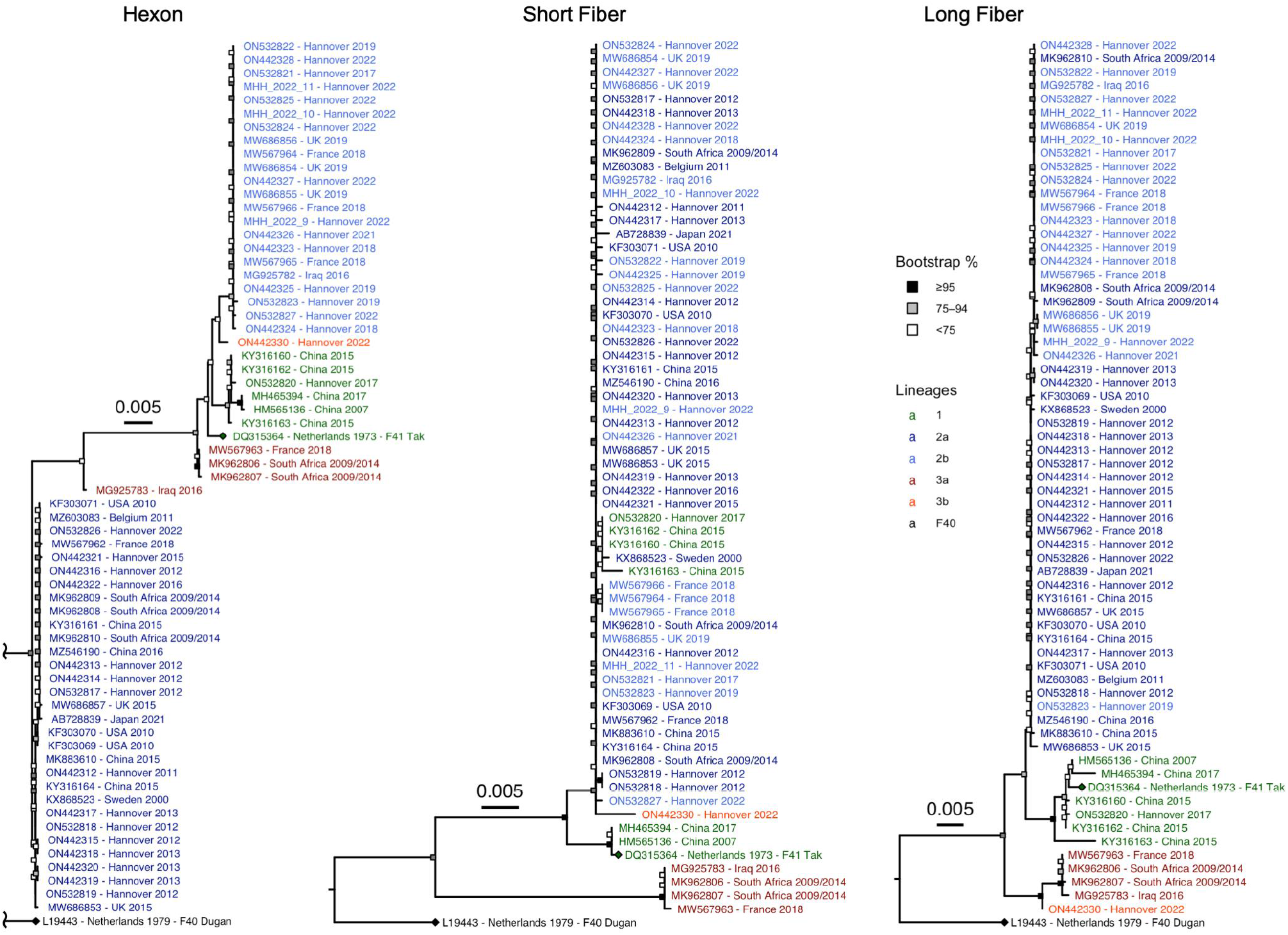
Phylogenetic trees of the hexon, short fiber, and long fiber genes of all 65 HAdV-F41 complete genomic sequences. The HAdV-F41 and -F40 prototype sequences are marked with a rhombus at the tip point. Bootstrap support values were binned into three categories (≥95 %, 75–94 %, <75 %) indicated by filled boxes at the node points. The lineage colouring from Figure 1 was retained. The total distance to the HAdV-F40 prototype was shortened in the hexon phylogeny due to the low genetic identity to the HAdV-F41 sequences (^~^81 %).

Other hotspots of evolution were in gene regions E3 and E4, coding for non-structural proteins (Figure 3). Lineages 3 had the lowest average sequence identity (93.6 %) compared to the prototype sequence in the E3 region, even lower than the sequence identity between HAdV-F41 prototype ‘Tak’ and HAdV-F40 prototype ‘Dugan’ (98.3 %). Sixty-four of 156 SNPs were non-synonymous, but the BUSTED algorithm did not confirm positive selection in any of the five E3 open reading frames (*p*=0.102 to *p*=0.5). In the E4 region, lineage 1 had 62 SNPs compared to the HAdV-F41 consensus sequence, with 23 being non-synonymous. Twenty-two of these were located in the E4 ORF4 and ORF6; however, no evidence for positive selection was found by BUSTED (*p*=0.5).

**Figure 3:**
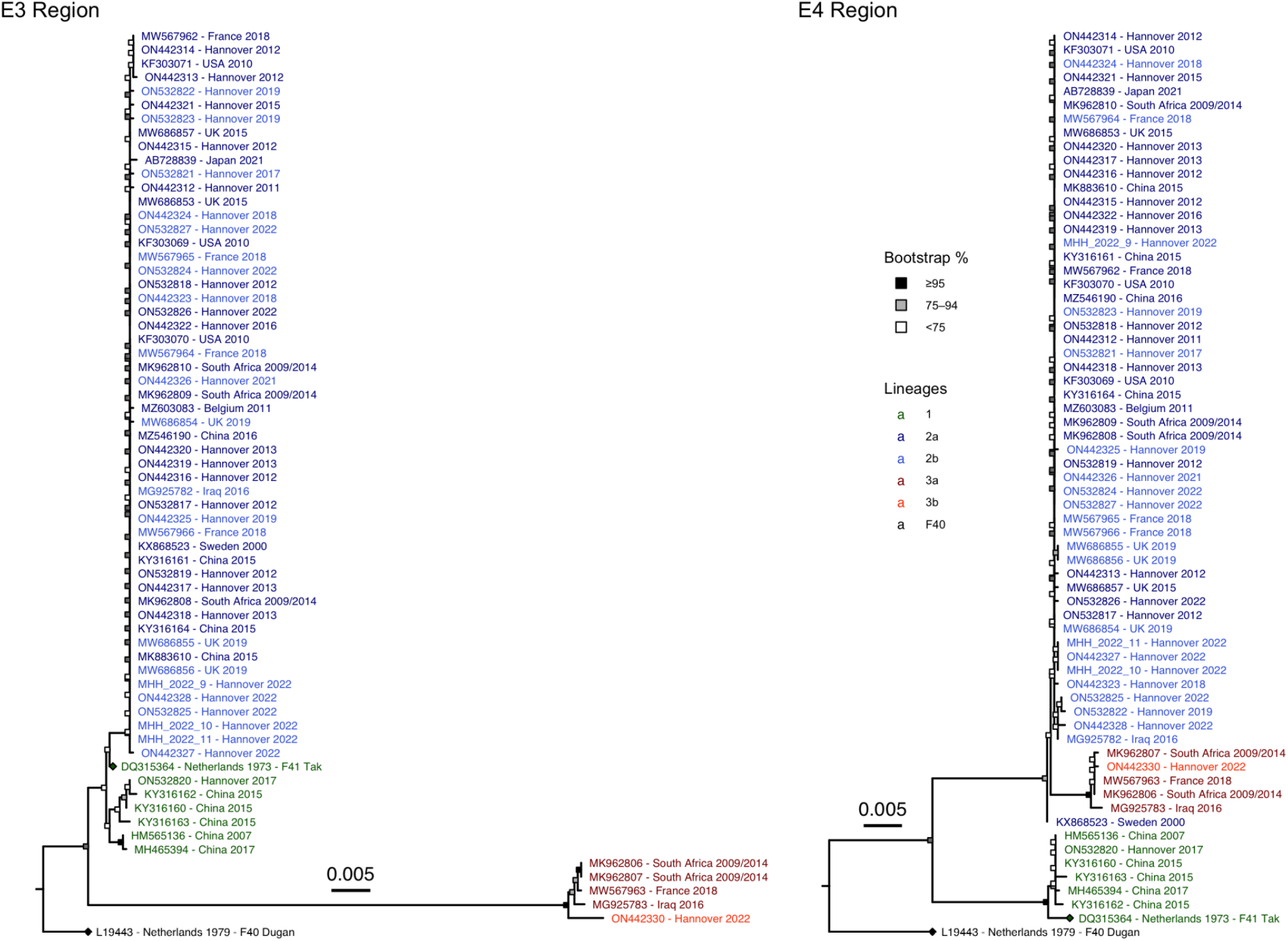
Phylogenetic trees of the E3 and E4 gene regions of all 65 HAdV-F41 complete genomic sequences. The HAdV-F41 and -F40 prototype sequences are marked with a rhombus at the tip point. Bootstrap support values were binned into three categories (≥95 %, 75–94 %, <75 %) indicated by filled boxes at the node points. The lineage colouring from Figure 1 was retained.

### Interlineage recombination

Intertypic recombination between the two HAdV species F types 40 and 41 was not observed in the phylogeny of HAdV-F41 lineages. However, the short fiber gene region of lineage 3b was phylogenetically linked to lineage 2, and the recombinant origin of this region was confirmed by bootscanning (Figure 4). The genome regions around the short fiber – E3, long fiber, and E4 – are phylogenetically clearly related to lineage 3a, while the genomes from the 5’-end to the E3 region were too similar between lineages 2 and 3a to show clear attribution in the BootScan.

**Figure 4:**
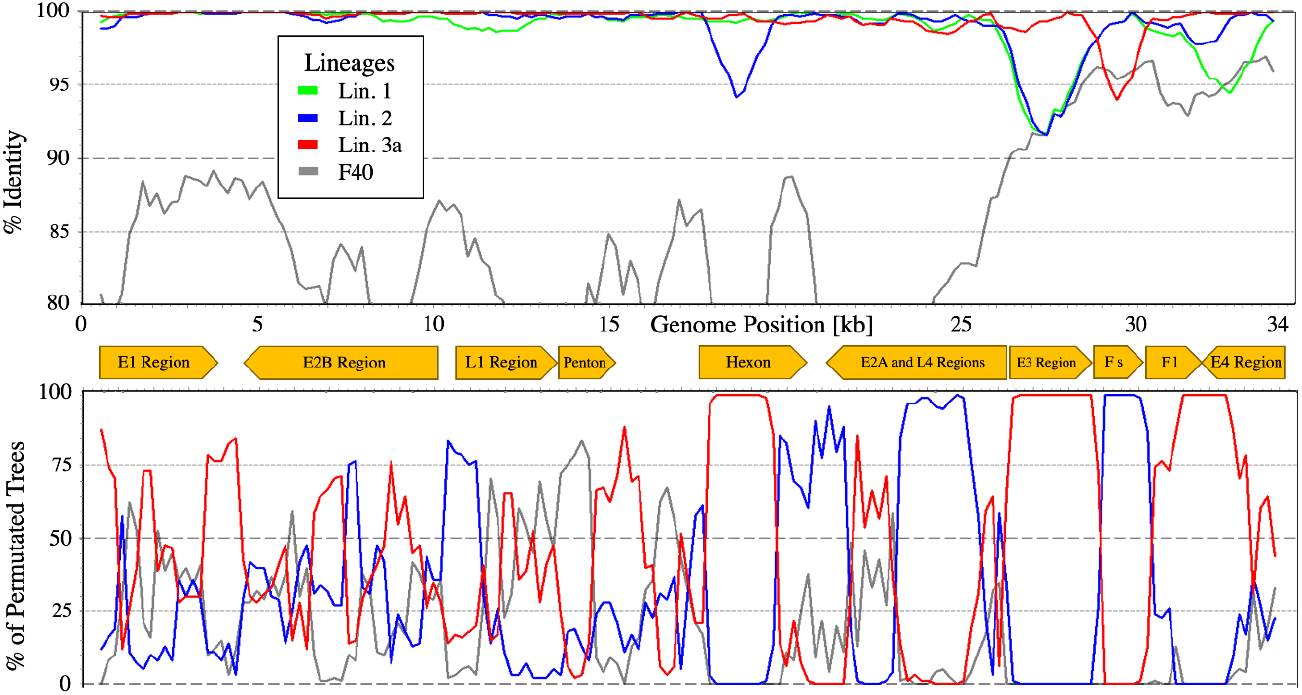
HAdV-F41 lineage 3b BootScan. BootScan plot of the lineage 3b genome with the consensus genomes of lineages 1, 2, and 3a, and HAdV-F40 as the outgroup.

Since no further interlineage recombinations were observed in the phylogeny of circulating HAdV-F41 strains, recombination was probably not a significant driver of the evolution of HAdV-F41 lineages.

## Discussion

HAdV-F41 was recently associated with severe hepatitis in children of about six months to two years of age, however, its aetiological significance has not yet been proven (Baker et al., 2022; Marsh et al., 2022; UK Health Security Agency, 2022b). These children had low, cell-associated virus loads (<10^5^ c/ml) in peripheral blood (Marsh et al., 2022), whereas typical adenovirus hepatitis causes high virus loads (usually >10^8^ c/ml) in plasma (Ronan et al., 2014; Schaberg et al., 2017). Moreover, adenovirus hepatitis is associated with life-threatening disseminated disease in severely immunosuppressed patients, e. g. haematopoietic stem cell transplant recipients (Forstmeyer et al., 2008; Lion, 2014; Onda et al., 2021). However, these hepatitis cases are rather caused by HAdV types of species C whereas HAdV-F41 infection remained limited to the gastrointestinal tract even in haematopoietic stem cell transplant recipients (Mattner et al., 2008; Lefeuvre et al., 2021). In the recent cases of severe hepatitis in children, detection of adenovirus antigens or viral particles failed in explanted liver specimens (Baker et al., 2022), whereas in typical adenovirus hepatitis, these can be found in abundance (Forstmeyer et al., 2008; Onda et al., 2021). Nevertheless, the emergence of a novel strain of HAdV-F41 could be suspected, which may be indirectly causing hepatitis. Unfortunately, recovery of high-quality HAdV-F41 genomes from hepatitis-affected children failed due to low virus loads (UK Health Security Agency, 2022b). HAdV-F41 has been highly endemic as a gastroenteritis pathogen for decades, but no cases were found in our collection between September 2019 and December 2021, probably due to hygiene measures implemented in Europe during the COVID-19 pandemic (Brauner et al., 2021; Marsh et al., 2022). An association of HAdV-F41 with hepatitis had not been observed previously but was described in the UK and several other countries during the re-emergence. Therefore, we generated complete genomic sequences of re-emergent HAdV-F41 clinical isolates (from 2021 and 2022) and archival isolates from 2011 to 2019. All these originated from gastroenteritis cases as hepatitis cases were unavailable to us. Generated data could either detect a novel strain during re-emergence or be compared to partial HAdV sequences generated from recently described hepatitis patients.

The vast majority of our complete genomic sequences, as well as available GenBank sequences, were clustered as lineage 2 (see Figure 1). This lineage – together with lineage 1 containing the prototype strain ‘Tak’ from 1973 – probably represented the abundant gastroenteritis strains encountered worldwide over several decades. Sublineage 2a may represent a partial immune-escape variant due to its mutations in the neutralisation determinant ε. Lineage 3 is characterised by a 15-amino acid deletion of the 15th repeat in the long fiber shaft, somewhat similar to the deletion of the 14th repeat in the long fiber shaft of HAdV-F40 (Kidd et al., 1990). Moreover, four amino acid substitutions were located in the long fiber knob, potentially affecting the binding to the cellular receptor. Furthermore, the E3 region of lineage 3 was highly divergent from both enteric species HAdV-F types 40 and 41 (lineages 1 and 2). Most E3 proteins exhibit immunomodulatory functions and can thus contribute to the virulence of an HAdV strain (Windheim et al., 2004).

Only sublineage 3a was significantly divergent in the short fiber, with four amino acid substitutions in the knob region. As the short fiber knob of HAdV-F41 was recently reported to bind to heparan sulfate, potentially restricting the cellular tropism, these mutations in sublineage 3a might lead to a circumvention of this restriction (Rajan et al., 2021). Surprisingly, the short fiber gene of sublineage 3b was found to have a recombinant phylogeny derived from lineage 2. Only a single sublineage 3b strain (from February 2022) was present in 65 complete genomic HAdV-F41 sequences, thus sublineage 3b may be considered as a novel recombinant strain. However, *in silico* restriction fragment length polymorphism (RFLP) analysis of the only sublineage 3b sequence revealed a genome type D6 (see Supplementary Table 2), which was already isolated in 1980 in the Netherlands and had the 15-amino acid deletion in the long fiber shaft (van der Avoort et al., 1989; Kidd et al., 1990). Identical RFLP patterns only elucidate the cutting sites of the ten used restriction enzymes and thus do not preclude significant divergence in other genome regions, which may have evolved recently and may influence tropism and virulence. Sublineage 3a genomes, on the other hand, were first identified by complete genomic sequencing of two samples from South Africa originating between 2009 and 2014 (MK962806 and MK962807). Furthermore, sublineage 3a genomes were found in Iraq (MG925783) and France (MW567963) as late as 2018 (Lefeuvre et al., 2021), but not in the present sequencing effort. The latter sequence originated from an HSCT patient and detailed virus load data were published (patient B in Lefeuvre et al., 2021). Interestingly, virus loads in stool were rather low (about 10^6^ c/ml) compared to other patients and DNAaemia appeared rather late and not in parallel to the peak virus load in stool as in other patients. Suggestively, this is somewhat similar to the DNAaemia pattern in the recently described hepatitis cases (Baker et al., 2022; UK Health Security Agency, 2022b, 2022a), but the highly immunocompromised patient B did not suffer from hepatitis. Perhaps an immune-mediated pathomechanism of liver injury, which is triggered by HAdV-F41 lineage 3 replication in other body sites (e.g. lymphoid tissue), can be suspected because severe hepatitis cases in children were associated with higher HAdV loads in blood (UK Health Security Agency, 2022a). Another speculative pathomechanism could be co-infection with adeno-associated virus 2 (AAV2), a dependoparvovirus, which may replicate in HAdV-F41–infected cells in other body sites than the liver. AAV2 DNAaemia was found in hepatitis patients as well as AAV2 DNA in the liver, in contrast to HAdV-F41 DNA (UK Health Security Agency, 2022b). AAV2 may perhaps cause an abortive infection of liver cells and thus liver cell injury or trigger an immunopathology against AAV2 antigens in liver cells.

In conclusion, the aetiology and pathomechanism of hepatitis cases in children associated with HAdV-F41 infections remain obscure and future research on these topics is urgently required.

## Author contributions

JG and AKC analysed and interpreted the data and drafted the manuscript. LS conducted the sequencing. AH designed and supervised the study, interpreted data, drafted and reviewed the manuscript. All authors contributed to the article and approved the submitted version.

## Acknowledgements

We thank Jenny Witthuhn and Sandra Flucht for their technical assistance.

## Data Availability

HAdV-F41 complete genomic sequences generated for this study are available at GenBank accession numbers (ON442312–ON442330, ON532817–ON532827). Accession numbers for pre-existing GenBank sequences used in this study are shown in the phylogenetic trees (Figures 1, 2, and 3).

## References

Albinsson, B., and Kidd, A. H. (1999). Adenovirus type 41 lacks an RGD alpha(v)-integrin binding motif on the penton base and undergoes delayed uptake in A549 cells. Virus Res 64, 125–136. doi: 10.1016/s0168-1702(99)00087-8.

Aoki, K., Benkő, M., Davison, A. J., Echavarria, M., Erdman, D. D., Harrach, B., et al. (2011). Toward an integrated human adenovirus designation system that utilizes molecular and serological data and serves both clinical and fundamental virology. J Virol 85, 5703–5704. doi: 10.1128/JVI.00491-11.

Baker, J. M., Buchfellner, M., Britt, W., Sanchez, V., Potter, J. L., Ingram, L. A., et al. (2022). Acute Hepatitis and Adenovirus Infection Among Children — Alabama, October 2021–February 2022. MMWR Morb. Mortal. Wkly. Rep. 71, 638–640. doi: 10.15585/mmwr.mm7118e1.

Brauner, J. M., Mindermann, S., Sharma, M., Johnston, D., Salvatier, J., Gavenčiak, T., et al. (2021). Inferring the effectiveness of government interventions against COVID-19. Science 371, eabd9338. doi: 10.1126/science.abd9338.

Davison, A. J., Benkő, M., and Harrach, B. (2003). Genetic content and evolution of adenoviruses. Journal of General Virology 84, 2895–2908. doi: 10.1099/vir.0.19497-0.

de Jong, J. C., Wigand, R., Kidd, A. H., Wadell, G., Kapsenberg, J. G., Muzerie, C. J., et al. (1983). Candidate adenoviruses 40 and 41: fastidious adenoviruses from human infant stool. J Med Virol 11, 215–231. doi: 10.1002/jmv.1890110305.

Dhingra, A., Hage, E., Ganzenmueller, T., Böttcher, S., Hofmann, J., Hamprecht, K., et al. (2019). Molecular Evolution of Human Adenovirus (HAdV) Species C. Sci Rep 9, 1039. doi: 10.1038/s41598-018-37249-4.

Forstmeyer, D., Henke-Gendo, C., Bröcker, V., Wildner, O., and Heim, A. (2008). Quantitative temporal and spatial distribution of adenovirus type 2 correlates with disease manifestations and organ failure during disseminated infection. J. Med. Virol. 80, 294–297. doi: 10.1002/jmv.21071.

Gonçalves, G., Gouveia, E., Mesquita, J. R., Almeida, A., Ribeiro, A., Rocha-Pereira, J., et al. (2011). Outbreak of acute gastroenteritis caused by adenovirus type 41 in a kindergarten. Epidemiol Infect 139, 1672–1675. doi: 10.1017/S0950268810002803.

Hilleman, M. R., and Werner, J. H. (1954). Recovery of new agent from patients with acute respiratory illness. Proc Soc Exp Biol Med 85, 183–188. doi: 10.3181/00379727-85-20825.

Katoh, K., and Standley, D. M. (2013). MAFFT multiple sequence alignment software version 7: improvements in performance and usability. Mol. Biol. Evol. 30, 772–780. doi: 10.1093/molbev/mst010.

Kidd, A. H., Erasmus, M. J., and Tiemessen, C. T. (1990). Fiber sequence heterogeneity in subgroup F adenoviruses. Virology 179, 139–150. doi: 10.1016/0042-6822(90)90283-W.

Lee, B., Damon, C. F., and Platts-Mills, J. A. (2020). Pediatric acute gastroenteritis due to adenovirus 40/41 in low- and middle-income countries. Curr Opin Infect Dis 33, 398–403. doi: 10.1097/QCO.0000000000000663.

Lefeuvre, C., Salmona, M., Feghoul, L., Ranger, N., Mercier-Delarue, S., Nizery, L., et al. (2021). Deciphering an Adenovirus F41 Outbreak in Pediatric Hematopoietic Stem Cell Transplant Recipients by Whole-Genome Sequencing. J Clin Microbiol 59, e03148–20. doi: 10.1128/JCM.03148-20.

Lion, T. (2014). Adenovirus Infections in Immunocompetent and Immunocompromised Patients. Clinical Microbiology Reviews 27, 441–462. doi: 10.1128/CMR.00116-13.

Lole, K. S., Bollinger, R. C., Paranjape, R. S., Gadkari, D., Kulkarni, S. S., Novak, N. G., et al. (1999). Full-length human immunodeficiency virus type 1 genomes from subtype C-infected seroconverters in India, with evidence of intersubtype recombination. J Virol 73, 152–160. doi: 10.1128/JVI.73.1.152-160.1999.

Marsh, K., Tayler, R., Pollock, L., Roy, K., Lakha, F., Ho, A., et al. (2022). Investigation into cases of hepatitis of unknown aetiology among young children, Scotland, 1 January 2022 to 12 April 2022. Eurosurveillance 27. doi: 10.2807/1560-7917.ES.2022.27.15.2200318.

Mattner, F., Sykora, K.-W., Meissner, B., and Heim, A. (2008). An adenovirus type F41 outbreak in a pediatric bone marrow transplant unit: analysis of clinical impact and preventive strategies. Pediatr Infect Dis J 27, 419–424. doi: 10.1097/INF.0b013e3181658c46.

Murrell, B., Weaver, S., Smith, M. D., Wertheim, J. O., Murrell, S., Aylward, A., et al. (2015). Gene-Wide Identification of Episodic Selection. Molecular Biology and Evolution 32, 1365–1371. doi: 10.1093/molbev/msv035.

Onda, Y., Kanda, J., Sakamoto, S., Okada, M., Anzai, N., Umadome, H., et al. (2021). Detection of adenovirus hepatitis and acute liver failure in allogeneic hematopoietic stem cell transplant patients. Transpl. Infect. Dis. 23. doi: 10.1111/tid.13496.

Rajan, A., Palm, E., Trulsson, F., Mundigl, S., Becker, M., Persson, B. D., et al. (2021). Heparan Sulfate Is a Cellular Receptor for Enteric Human Adenoviruses. Viruses 13. doi: 10.3390/v13020298.

Robinson, C. M., Singh, G., Lee, J. Y., Dehghan, S., Rajaiya, J., Liu, E. B., et al. (2013). Molecular evolution of human adenoviruses. Scientific Reports 3, 1812. doi: 10.1038/srep01812.

Roelvink, P. W., Lizonova, A., Lee, J. G. M., Li, Y., Bergelson, J. M., Finberg, R. W., et al. (1998). The Coxsackievirus-Adenovirus Receptor Protein Can Function as a Cellular Attachment Protein for Adenovirus Serotypes from Subgroups A, C, D, E, and F. J Virol 72, 7909–7915. doi: 10.1128/JVI.72.10.7909-7915.1998.

Ronan, B. A., Agrwal, N., Carey, E. J., De Petris, G., Kusne, S., Seville, M. T., et al. (2014). Fulminant hepatitis due to human adenovirus. Infection 42, 105–111. doi: 10.1007/s15010-013-0527-7.

Ronquist, F., Teslenko, M., van der Mark, P., Ayres, D. L., Darling, A., Höhna, S., et al. (2012). MrBayes 3.2: efficient Bayesian phylogenetic inference and model choice across a large model space. Syst Biol 61, 539–542. doi: 10.1093/sysbio/sys029.

Rowe, W. P., Huebner, R. J., Gilmore, L. K., Parrott, R. H., and Ward, T. G. (1953). Isolation of a Cytopathogenic Agent from Human Adenoids Undergoing Spontaneous Degeneration in Tissue Culture. Proceedings of the Society for Experimental Biology and Medicine 84, 570–573. doi: 10.3181/00379727-84-20714.

Sagulenko, P., Puller, V., and Neher, R. A. (2018). TreeTime: Maximum-likelihood phylodynamic analysis. Virus Evolution 4, vex042. doi: 10.1093/ve/vex042.

Schaberg, K. B., Kambham, N., Sibley, R. K., and Higgins, J. P. T. (2017). Adenovirus Hepatitis: Clinicopathologic Analysis of 12 Consecutive Cases From a Single Institution. American Journal of Surgical Pathology 41, 810–819. doi: 10.1097/PAS.0000000000000834.

Seto, D., Chodosh, J., Brister, J. R., Jones, M. S., and Community, M. of the A. R. (2011). Using the Whole-Genome Sequence To Characterize and Name Human Adenoviruses. Journal of Virology 85, 5701–5702. doi: 10.1128/JVI.00354-11.

Slatter, M. A., Read, S., Taylor, C. E., Crooks, B. N. A., Abinun, M., Flood, T. J., et al. (2005). Adenovirus Type F Subtype 41 Causing Disseminated Disease following Bone Marrow Transplantation for Immunodeficiency. J Clin Microbiol 43, 1462–1464. doi: 10.1128/JCM.43.3.1462-1464.2005.

Stamatakis, A. (2014). RAxML version 8: a tool for phylogenetic analysis and post-analysis of large phylogenies. Bioinformatics 30, 1312–1313. doi: 10.1093/bioinformatics/btu033.

Tahmasebi, R., Luchs, A., Tardy, K., Hefford, P. M., Tinker, R. J., Eilami, O., et al. (2020). Viral gastroenteritis in Tocantins, Brazil: characterizing the diversity of human adenovirus F through next-generation sequencing and bioinformatics. J Gen Virol 101, 1280–1288. doi: 10.1099/jgv.0.001500.

UK Health Security Agency (2022a). Acute hepatitis of unknown aetiology: Technical briefing 1. Available at: https://assets.publishing.service.gov.uk/government/uploads/system/uploads/attachment_data/file/1071198/acute-hepatitis-technical-briefing-1_4_.pdf.

UK Health Security Agency (2022b). Acute hepatitis of unknown aetiology: Technical briefing 2. Available at: https://assets.publishing.service.gov.uk/government/uploads/system/uploads/attachment_data/file/1073704/acute-hepatitis-technical-briefing-2.pdf.

van der Avoort, H. G., Wermenbol, A. G., Zomerdijk, T. P., Kleijne, J. A., van Asten, J. A., Jensma, P., et al. (1989). Characterization of fastidious adenovirus types 40 and 41 by DNA restriction enzyme analysis and by neutralizing monoclonal antibodies. Virus Res 12, 139–157. doi: 10.1016/0168-1702(89)90060-9.

Walker, B. J., Abeel, T., Shea, T., Priest, M., Abouelliel, A., Sakthikumar, S., et al. (2014). Pilon: An Integrated Tool for Comprehensive Microbial Variant Detection and Genome Assembly Improvement. PLOS ONE 9, e112963. doi: 10.1371/journal.pone.0112963.

Weaver, S., Shank, S. D., Spielman, S. J., Li, M., Muse, S. V., and Kosakovsky Pond, S. L. (2018). Datamonkey 2.0: A Modern Web Application for Characterizing Selective and Other Evolutionary Processes. Molecular Biology and Evolution 35, 773–777. doi: 10.1093/molbev/msx335.

Wickham, H. (2016). ggplot2: Elegant graphics for data analysis. Springer-Verlag New York Available at: https://ggplot2.tidyverse.org.

Wickham, T. J., Mathias, P., Cheresh, D. A., and Nemerow, G. R. (1993). Integrins alpha v beta 3 and alpha v beta 5 promote adenovirus internalization but not virus attachment. Cell 73, 309–319. doi: 10.1016/0092-8674(93)90231-e.

Windheim, M., Hilgendorf, A., and Burgert, H.-G. (2004). “Immune Evasion by Adenovirus E3 Proteins: Exploitation of Intracellular Trafficking Pathways,” in Adenoviruses: Model and Vectors in Virus-Host Interactions Current Topics in Microbiology and Immunology., eds. W. Doerfler and P. Böhm (Berlin, Heidelberg: Springer Berlin Heidelberg), 29–85. doi: 10.1007/978-3-662-05599-1_2.

Yu, G. (2020). Using ggtree to Visualize Data on Tree-Like Structures. Current Protocols in Bioinformatics 69, e96. doi: 10.1002/cpbi.96.

