## Supplementary Material for "Molecular Phylogeny of human adenovirus type 41 lineages"

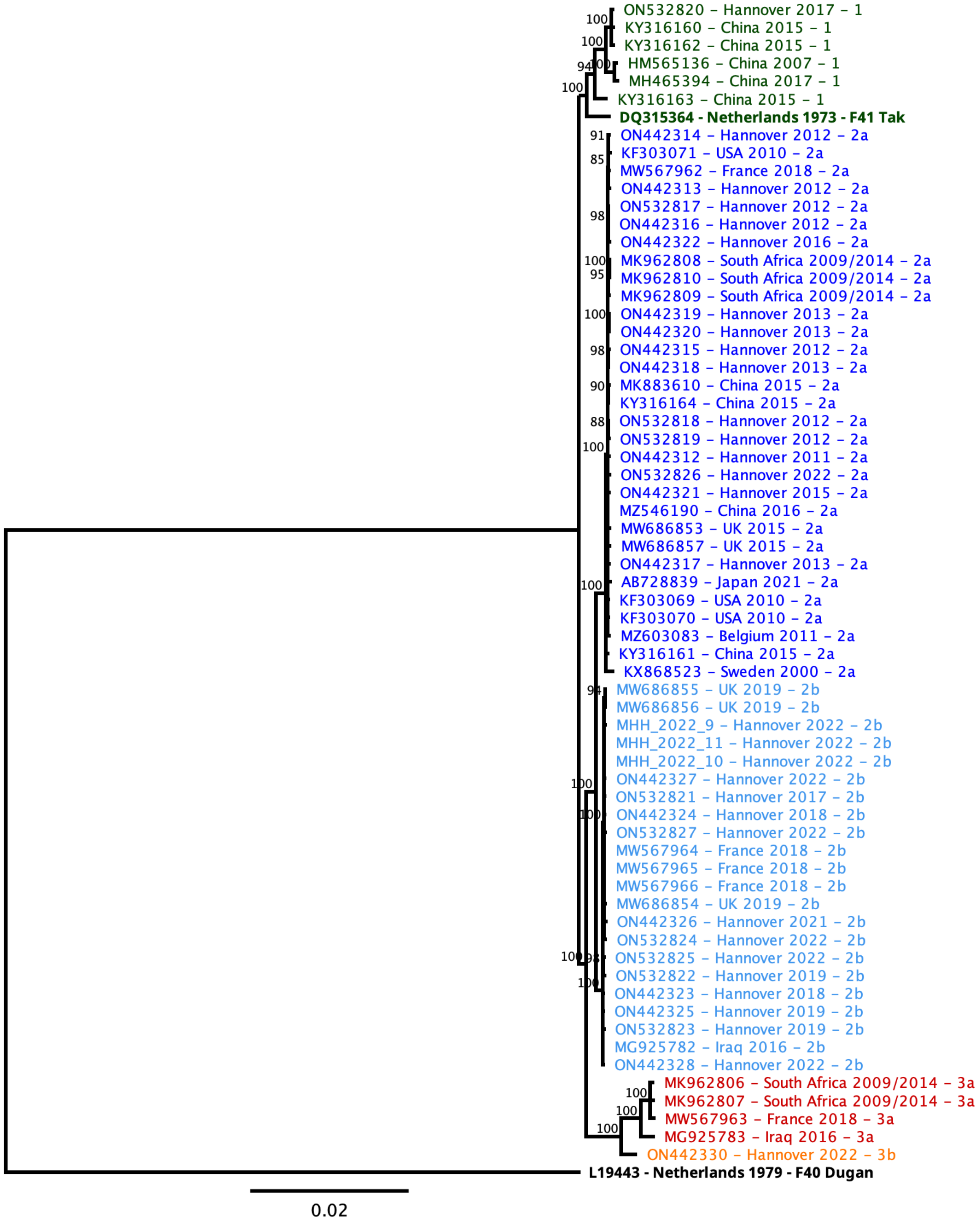


**Supplementary Figure 1:** Neighbour-joining phylogeny of all 65 HAdV-F41 genomes with the HAdV-F40 prototype as the outgroup. The two prototype sequences (HAdV-F41 DQ315364, HAdV-F40 L19443) are marked by a bold label. The lineage number from Figure 1 is appended to the tip label and lineage colouring has been retained. Support values from 1000 bootstrap replicates are shown at the node points.

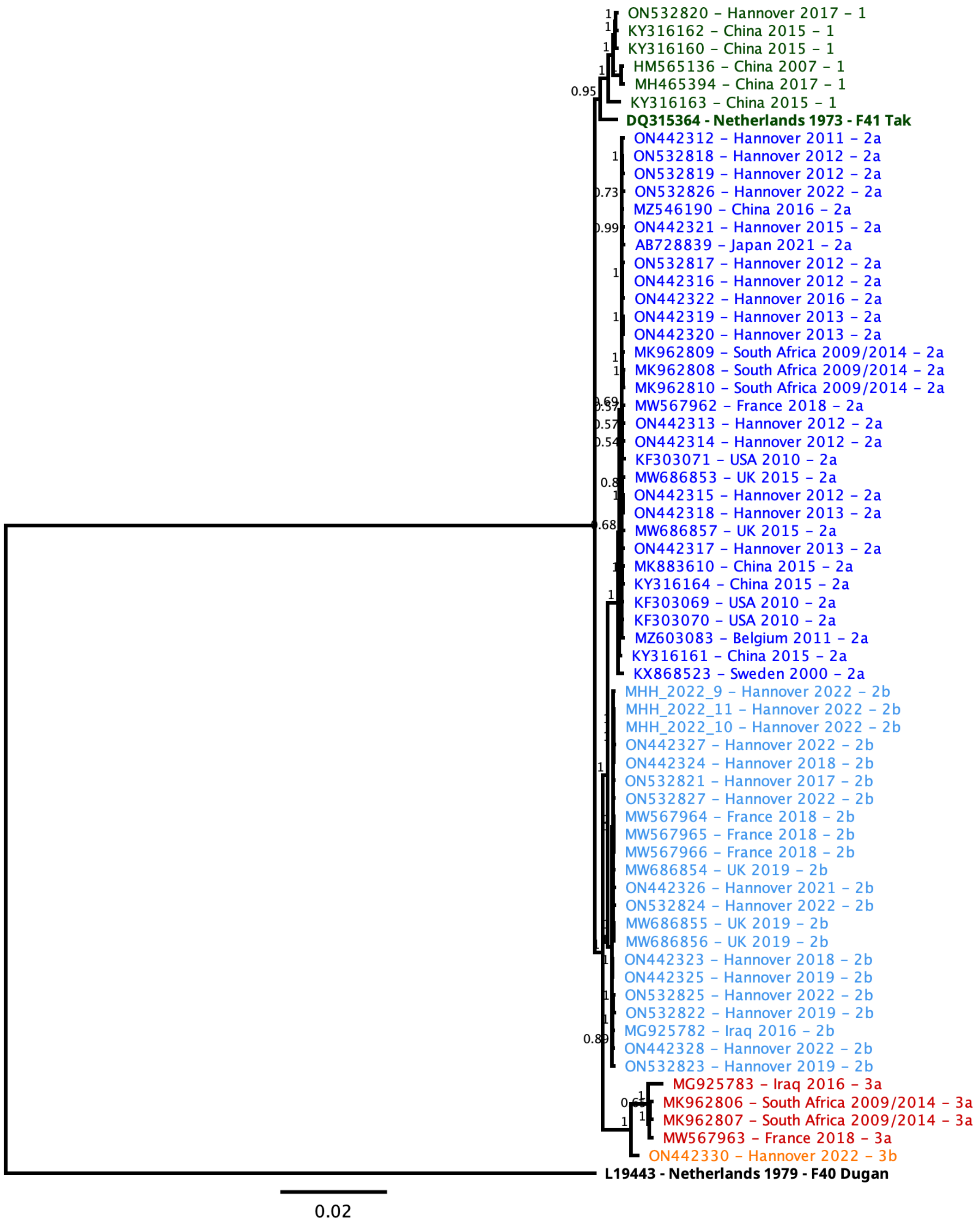


**Supplementary Figure 2:** Bayesian phylogeny of all 65 HAdV-F41 genomes with the HAdV-F40 prototype as the outgroup. The two prototype sequences (HAdV-F41 DQ315364, HAdV-F40 L19443) are marked by a bold label. The lineage number from Figure 1 is appended to the tip label and lineage colouring has been retained. Posterior probabilities are shown at the node points.

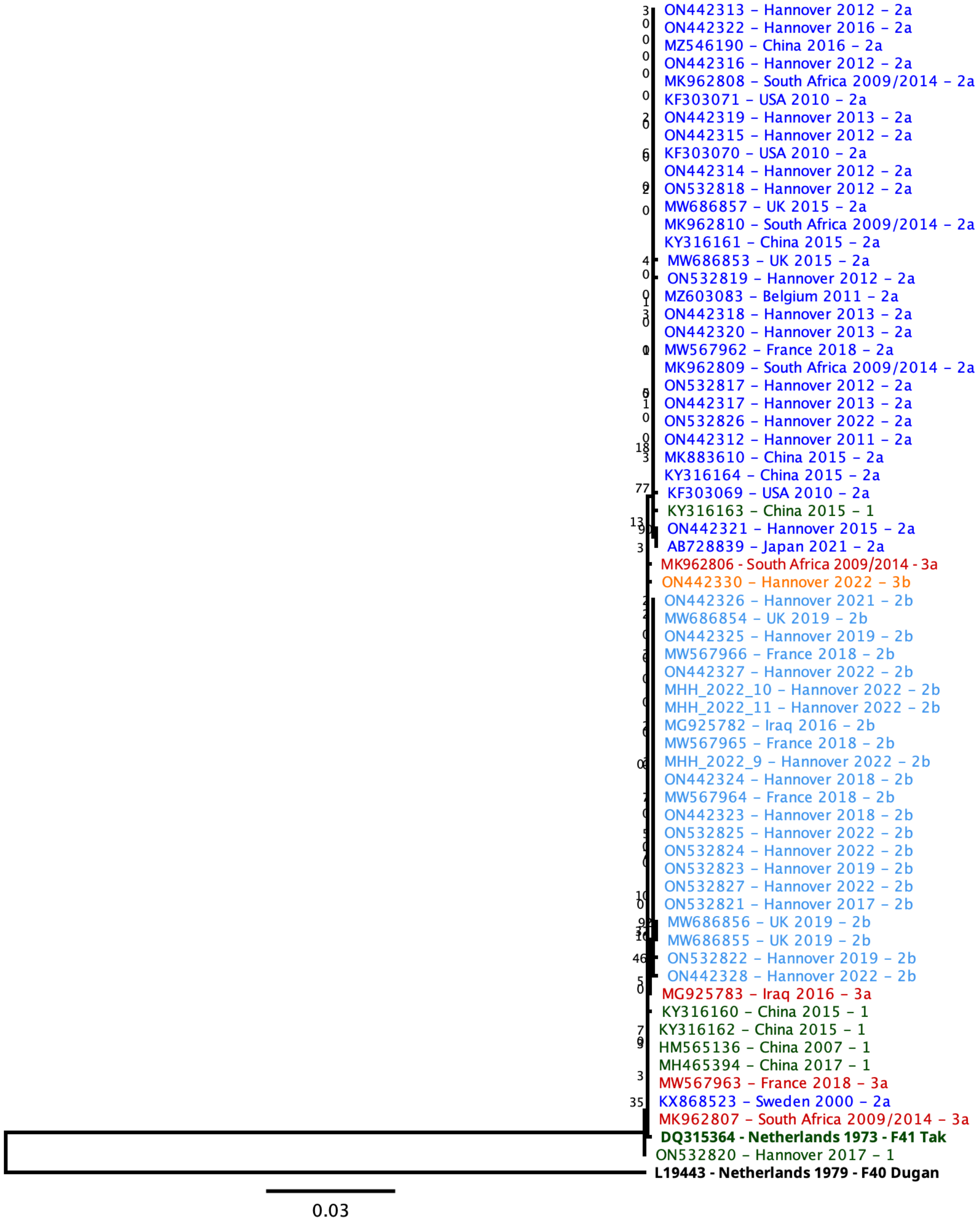


**Supplementary Figure 3:** Maximum-likelihood phylogeny of the penton base gene from all 65 HAdV-F41 genomes with the HAdV-F40 prototype as the outgroup. The two prototype sequences (HAdV-F41 DQ315364, HAdV-F40 L19443) are marked by a bold label. The lineage number from Figure 1 is appended to the tip label and lineage colouring has been retained. Bootstrap support values are shown at the node points.

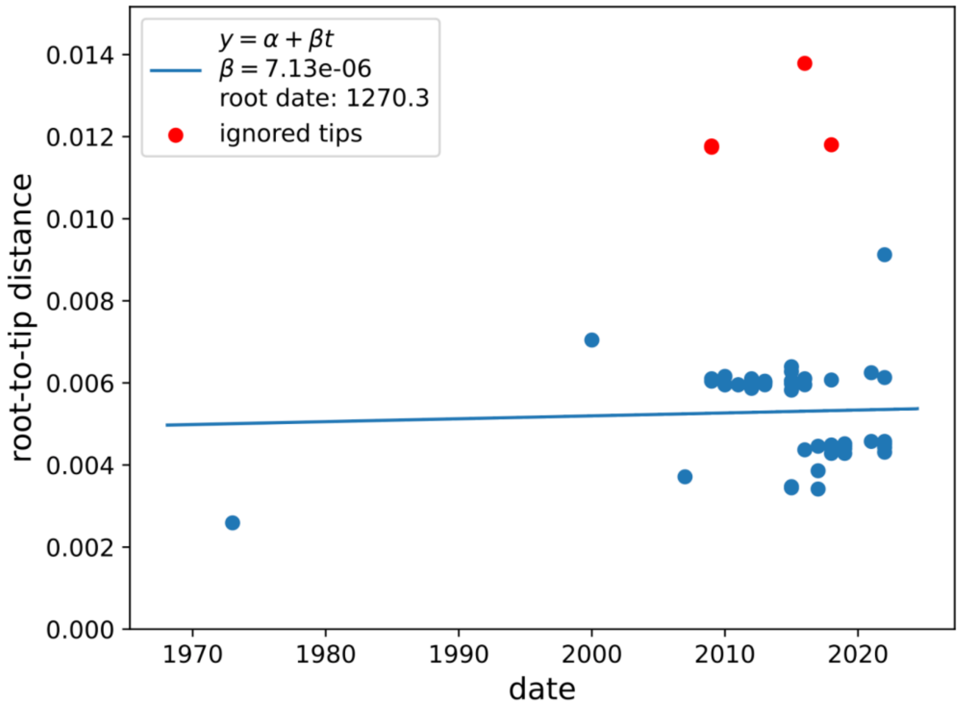


**Supplementary Figure 4:** Scatterplot of root-to-tip distance in the CGS phylogenetic tree versus year of isolation ($R^2$ = 0.00). Correlation and plot generated using TreeTime.

**Supplementary Table 1:** BUSTED results from datamonkey.org

| CDS | Model | log L | Param No. | AICc | CV(SRV) | Branch set | ω1 | ω2 | ω3 | LRT p-value | Conclusion |
|---|---|---|---|---|---|---|---|---|---|---|---|
| **Hexon** | Unconstrained | -4589.3 | 71 | 9321.1 | 3.177 | Test | 0.04(1.26%) | 0.04(98.28%) | 14.34(0.46%) | 0.087 | No evidence |
| | Constrained | -4591.1 | 70 | 9322.6 | 3.019 | Test | 0.03(4.96%) | 0.03(90.04%) | 1.00 (5.01%) | | |
| **Short Fiber** | Unconstrained | -2029.0 | 59 | 4177.0 | 0.343 | Test | 0.00(64.45%) | 0.00(22.03%) | 1.64(13.52%) | 0.444 | No evidence |
| | Constrained | -2029.1 | 58 | 4175.2 | 0.482 | Test | 0.00(48.18%) | 0.00(31.16%) | 1.00(20.66%) | | |
| **Short Fiber Knob** | Unconstrained | -687.4 | 39 | 1455.4 | 0.000 | Test | 0.24 (88.72%) | 0.65 (11.28%) | 4.22 (0.00%) | 0.500 | No evidence |
| **Long Fiber** | Unconstrained | -2772.4 | 63 | 5671.5 | 1.328 | Test | 0.00 (3.23%) | 0.09(96.46%) | 344.45(0.31%) | **0.000** | **Found evidence** |
| | Constrained | -2780.7 | 62 | 5686.1 | 1.583 | Test | 0.00(66.27%) | 0.00(15.19%) | 1.00 (18.54%) | | |
| **Long Fiber Knob** | Unconstrained | -832.7 | 39 | 1745.6 | 0.742 | Test | 0.00(79.12%) | 0.00(12.26%) | 3.21 (8.63%) | 0.326 | No evidence |
| | Constrained | -833.1 | 38 | 1744.3 | 1.008 | Test | 0.00(69.17%) | 0.00 (8.61%) | 1.00(22.22%) | | |
| **E3 10.1K** | Unconstrained | -432.2 | 35 | 938.6 | 0.894 | Test | 0.06 (0.00%) | 0.09 (100.00%) | 1.16 (0.00%) | 0.500 | No evidence |
| **E3 14.5K** | Unconstrained | -551.1 | 41 | 1187.4 | 0.733 | Test | 0.22 (0.00%) | 0.22 (100.00%) | 3.98 (0.00%) | 0.500 | No evidence |
| **E3 14.7K** | Unconstrained | -543.6 | 33 | 1156.5 | 2.812 | Test | 0.00(80.47%) | 0.00(13.76%) | 39.78(5.77%) | 0.102 | No evidence |
| | Constrained | -545.2 | 32 | 1157.4 | 2.612 | Test | 0.00(67.19%) | 0.00(10.19%) | 1.00(22.61%) | | |
| **E3 19.4K** | Unconstrained | -888.2 | 39 | 1856.4 | 1.215 | Test | 0.00 (46.70%) | 0.84 (53.30%) | 1.30 (0.00%) | 0.500 | No evidence |
| **E3 31.6K** | Unconstrained | -1503.8 | 43 | 3094.8 | 1.253 | Test | 0.15(54.47%) | 0.15(20.16%) | 2.70(25.38%) | 0.342 | No evidence |
| | Constrained | -1504.1 | 42 | 3093.5 | 0.657 | Test | 0.00(31.08%) | 1.00(14.04%) | 1.00(54.88%) | | |
| **E4 ORF4** | Unconstrained | -602.9 | 39 | 1286.9 | 28.387 | Test | 0.31 (0.00%) | 0.38 (100.00%) | 1.00 (0.00%) | 0.500 | No evidence |
| **E4 ORF6** | Unconstrained | -1472.9 | 49 | 3044.9 | 0.908 | Test | 0.19(100.00%) | 0.23(0.00%) | 5.17(0.00%) | 0.500 | No evidence |

**Supplementary Table 2:** RFLP enzyme pattern numbers from the *in silico* RFLP analysis using the ten restriction enzymes from van der Avoort et al., 1989. Black numbers indicate patterns described by van der Avoort et al., bold red numbers indicate novel patterns not described previously.

| Sample | BamHI | BglI | BstEII | EcoRI | HindIII | KpnI | PstI | SacI | SmaI | XhoI |
|---|---|---|---|---|---|---|---|---|---|---|
| **17358145 - 2017 - 1** | 2 | 3 | 2 | **7** | 1 | 1 | 1 | 1 | 2 | 2 |
| **DQ315364 - 1973 - 1** | 1 | 1 | 1 | 1 | 1 | 1 | 1 | 1 | 1 | 1 |
| **HM565136 - 2007 - 1** | 2 | 3 | 1 | **7** | 1 | 1 | 1 | **3** | 4 | 2 |
| **KY316160 - 2015 - 1** | 2 | 3 | 2 | **7** | 1 | 1 | 1 | 1 | 2 | 2 |
| **KY316162 - 2015 - 1** | 2 | 3 | 2 | **7** | 1 | 1 | 1 | 1 | 2 | 2 |
| **KY316163 - 2015 - 1** | 2 | 3 | 2 | **7** | 1 | 1 | 1 | 1 | 2 | 1 |
| **MH465394 - 2017 - 1** | 2 | 3 | 1 | **7** | 1 | 1 | **4** | 1 | 4 | 2 |
| **854751 - 2011 - 2a** | 2 | 1 | 2 | 1 | 5 | 1 | 1 | 1 | 2 | 1 |
| **882339 - 2012 - 2a** | 2 | 1 | 2 | 1 | 5 | 1 | 1 | 1 | 2 | 1 |
| **923682 - 2012 - 2a** | 2 | 1 | 2 | 1 | 5 | 1 | 1 | 1 | 2 | 1 |
| **928029 - 2012 - 2a** | 2 | 1 | 2 | 1 | 5 | 1 | 1 | 1 | 2 | 1 |
| **928030 - 2012 - 2a** | 2 | 1 | 2 | 1 | 5 | 1 | 1 | 1 | 2 | 1 |
| **928794 - 2012 - 2a** | 2 | 1 | 2 | 1 | 5 | 1 | 1 | 1 | 2 | 1 |
| **940923 - 2013 - 2a** | 2 | 1 | 2 | 1 | 5 | 1 | 1 | 1 | 2 | 1 |
| **940925 - 2012 - 2a** | 2 | 1 | 2 | 1 | 5 | 1 | 1 | 1 | 2 | 1 |
| **940927 - 2013 - 2a** | 2 | 1 | 2 | 1 | 5 | 1 | 1 | 1 | 2 | 1 |
| **1053305 - 2016 - 2a** | 2 | 1 | 2 | 1 | 5 | 1 | 1 | 1 | 2 | 1 |
| **12213478 - 2012 - 2a** | 2 | 1 | 2 | 1 | 5 | 1 | 1 | 1 | 2 | 1 |
| **12603313 - 2013 - 2a** | 2 | 1 | 2 | 1 | 5 | 1 | 1 | 1 | 2 | 1 |
| **12681552 - 2013 - 2a** | 2 | 1 | 2 | 1 | 5 | 1 | 1 | 1 | 2 | 1 |
| **15365373 - 2015 - 2a** | 2 | 1 | 2 | 1 | 5 | 1 | 1 | 1 | 2 | 1 |
| **20303866 - 2011 - 2a** | 2 | 1 | 2 | 1 | 5 | 1 | 1 | 1 | 2 | 1 |
| **60456594 - 2022 - 2a** | 2 | 1 | 2 | 1 | 5 | 1 | 1 | 1 | 2 | 1 |
| **AB728839 - 2021 - 2a** | 2 | 1 | 2 | 1 | 5 | 1 | 1 | 1 | 2 | 1 |
| **KF303069 - 2010 - 2a** | 2 | 1 | 2 | 1 | 5 | 1 | 1 | 1 | 2 | 1 |
| **KF303070 - 2010 - 2a** | 2 | 1 | 2 | 1 | 5 | 1 | 1 | 1 | 2 | 1 |
| **KF303071 - 2010 - 2a** | 2 | 1 | 2 | 1 | 5 | 1 | 1 | 1 | 2 | 1 |
| **KX868523 - 2000 - 2a** | 2 | 1 | 2 | 1 | 5 | 1 | 3 | 1 | 2 | 1 |
| **KY316161 - 2015 - 2a** | 2 | 1 | 2 | 1 | 5 | 1 | 1 | 1 | 2 | 1 |
| **KY316164 - 2015 - 2a** | 2 | 1 | 2 | 1 | 5 | 1 | 1 | 1 | 2 | 1 |
| **MK883610 - 2015 - 2a** | 2 | 1 | **6** | 1 | 5 | 1 | 1 | 1 | 2 | 1 |
| **MK962809 - 2009/2014 - 2a** | 2 | 1 | 2 | 4 | 5 | 1 | 1 | 1 | 2 | 1 |
| **MK962810 - 2009/2014 - 2a** | 2 | 1 | 2 | 1 | 5 | 1 | 1 | 1 | 2 | 1 |
| **MW567962 - 2018 - 2a** | 2 | 1 | 2 | 1 | 5 | 1 | 1 | 1 | 2 | 1 |
| **MW686853 - 2015 - 2a** | 2 | 1 | 2 | 1 | 5 | 1 | 1 | 1 | 2 | 1 |
| **MW686857 - 2015 - 2a** | 2 | 1 | 2 | 1 | 5 | 1 | 1 | 1 | 2 | 1 |
| **MZ546190 - 2016 - 2a** | 2 | 1 | 2 | 1 | 5 | 1 | 1 | 1 | 2 | 1 |
| **1119364 - 2018 - 2b** | 2 | 1 | 2 | 1 | 2 | 1 | 1 | 1 | 5 | 1 |

| | | | | | | | | | | |
|---|---|---|---|---|---|---|---|---|---|---|
| **1231469 - 2021 - 2b** | 2 | 1 | 2 | 1 | 2 | 1 | 1 | 1 | 5 | 1 |
| **1236866 - 2022 - 2b** | 2 | 1 | 2 | 1 | 2 | 1 | 1 | 1 | 5 | 1 |
| **17699366 - 2017 - 2b** | 2 | 1 | 2 | 1 | 2 | 1 | 1 | 1 | 5 | 1 |
| **60041744 - 2018 - 2b** | 2 | 1 | 2 | 1 | 2 | 1 | 1 | 1 | 5 | 1 |
| **60047233 - 2019 - 2b** | 2 | 1 | 2 | 1 | 2 | 1 | 1 | 1 | 5 | 1 |
| **60089894 - 2019 - 2b** | 2 | 1 | 2 | 1 | 2 | 1 | 1 | 1 | 5 | 1 |
| **60093021 - 2019 - 2b** | 2 | 1 | 2 | 1 | 2 | 1 | 1 | 1 | 5 | 1 |
| **60392103 - 2022 - 2b** | 2 | 1 | 2 | 1 | 2 | 1 | 1 | 1 | 5 | 1 |
| **60403125 - 2022 - 2b** | 2 | 1 | 2 | 1 | 2 | 1 | 1 | 1 | 5 | 1 |
| **60453790 - 2022 - 2b** | 2 | 1 | 2 | 1 | 2 | 1 | 1 | 1 | 5 | 1 |
| **60457773 - 2022 - 2b** | 2 | 1 | 2 | 1 | 2 | 1 | 1 | 1 | 5 | 1 |
| **MG925782 - 2016 - 2b** | 2 | 1 | 2 | 1 | 2 | 1 | 1 | 1 | 5 | 1 |
| **MW567964 - 2018 - 2b** | 2 | 1 | 2 | 1 | 2 | 1 | 1 | 1 | 5 | 1 |
| **MW567965 - 2018 - 2b** | 2 | 1 | 2 | 1 | 2 | 1 | 1 | 1 | 5 | 1 |
| **MW567966 - 2018 - 2b** | 2 | 1 | 2 | 1 | 2 | 1 | 1 | 1 | 5 | 1 |
| **MW686854 - 2019 - 2b** | 2 | 1 | 2 | 1 | 2 | 1 | 1 | 1 | 5 | 1 |
| **MW686855 - 2019 - 2b** | 2 | 1 | 2 | 1 | 2 | 1 | 1 | 1 | 5 | 1 |
| **MW686856 - 2019 - 2b** | 2 | 1 | 2 | 1 | 2 | 1 | 1 | 1 | 5 | 1 |
| **MG925783 - 2016 - 3a** | 2 | 1 | **5** | 4 | 6 | 2 | **5** | 1 | **10** | 1 |
| **MK962806 - 2009/2014 - 3a** | 2 | 1 | 4 | 4 | 4 | 2 | **5** | 1 | **9** | 1 |
| **MK962807 - 2009/2014 - 3a** | 2 | 1 | 4 | 4 | 4 | 2 | **5** | 1 | **9** | 1 |
| **MW567963 - 2018 - 3a** | 2 | 1 | 4 | 4 | 4 | 2 | **5** | 1 | **9** | 1 |
| **60425148 - 2022 - 3b** | 2 | 1 | 3 | 4 | 4 | 2 | 1 | 1 | 2 | 1 |
